# Neurobiology as Information Physics

**DOI:** 10.1101/060467

**Authors:** Sterling Street

## Abstract

This article reviews thermodynamic relationships in the brain in an attempt to consolidate current research in systems neuroscience. The present synthesis supports proposals that thermodynamic information in the brain can be quantified to an appreciable degree of objectivity, that many qualitative properties of information in systems of the brain can be inferred by observing changes in thermodynamic quantities, and that many features of the brain’s anatomy and architecture illustrate relatively simple information-energy relationships. The brain may provide a unique window into the relationship between energy and information.

## Introduction

That information is physical has been suggested by evidence since the founding of classical thermodynamics (J Gleick 2011; S Lloyd 2006). In recent years, Landauer’s principle (CH Bennett 2003; R Landauer 1996), which relates information-theoretic entropy to thermodynamic information, has been confirmed (JMR Parrondo et al. 2015), and the experimental demonstration of a form of information-energy equivalence (A Alfonso-Faus 2013) has verified that Maxwell’s demon cannot violate any known laws of thermodynamics (K Maruyama et al. 2009). The theoretical finding that entropy is conserved as event horizon area is leading to the resolution of the black hole information paradox (P Davies 2010; C Moskowitz 2015), and there is a fundamental relationship between information and the geometry of spacetime itself (R Bousso 2002; C Eling et al. 2006). Current formulations of quantum theory are revealing properties of physical information (Č Brukner and A Zeilinger 2003; S Lloyd 2006; V Vedral 2010; J Wheeler 1986), and information-interpretive attempts to show that gravity is quantized (JW Lee et al. 2013; L Smolin 2001) could even lead to the unification of quantum mechanics and the theories of relativity. Although similar approaches are increasingly influential in biology (JL England 2013; JC Flack 2014; ED Schneider and D Sagan 2005), “a formalization of the relationship between information and energy is currently lacking in neuroscience” (G Collell and J Fauquet 2015). The purpose of this article is to explore a few different sides of this relationship and, along the way, to suggest that many hypotheses and theories in neuroscience can be unified by the physics of information.

## Information bounds

> *“How can the events in space and time which take place within the spatial boundary of a living organism be accounted for by physics and chemistry?”*–
>
> — (E Schrödinger 1944, from KJ Friston 2013)

As a fundamental physical entity (S Lloyd 2015), information is not fully understood, and there is currently a significant amount of disagreement over different definitions of information and entropy in the literature (A Ben-Naim 2015; B Poirier 2014). In thermodynamics, however, information can be defined as a negation of thermodynamic entropy (C Beck 2009):

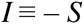

A bit of thermodynamic entropy represents the distinction between two alternative states in a physical system (JV Stone 2015). As a result, the total thermodynamic entropy of a system is proportional to the total number of distinguishable states contained in the system (JD Bekenstein 2001; JD Bekenstein 2007). Because thermodynamic entropy is potential information relative to an observer (S Lloyd 2006), and an observer in a physical system is a component of the system itself, the total thermodynamic entropy of a system includes the portion of entropy that is accessible to the observer as relative thermodynamic information (G Collell and J Fauquet 2015; J Wheeler 1989):

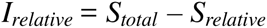

Since entropy in any physical system is finite (S Lloyd 2006; C Rovelli 2015), the total thermodynamic entropy of any system of the brain can be quantified by applying the traditional form of the universal (JD Bekenstein 1981, 1984, 2001, 2004, 2007) information-entropy bound:

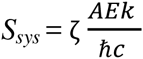

where *A* is area, *E* is energy including matter, ℏ is the reduced Planck constant, *c* is the speed of light, *k* is Boltzmann’s constant, and ζ is a factor such that 0 ≤ ζ ≤ 1

Setting this factor to 1 in order to quantify the total thermodynamic entropy of a system at a certain level of structure now allows us to quantifyζ thermodynamicζ information by partitioningζ the factor into a relative information component (*ζ*_*I*_ = 1 − *ζ*_*I*_) and a relative entropy component (*ζ*_*I*_ = 1 − *ζ*_*I*_),

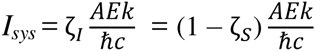

Because a maximal level of energy corresponds to a maximal level of thermodynamic information, and a minimal level of energy corresponds to a minimal level of thermodynamic information (TL Duncan and JS Semura 2004), any transitions between energy levels occur as transitions between informational extrema. So, in the event that information enters a system of the brain,

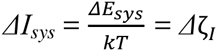

where *T* is temperature

and, in the event that information exits a system,

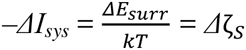

Various forms of these relationships, including information-entropy bounds, have been applied in neuroscience (G Collell and J Fauquet 2015; B Sengupta et al. 2013; B Sengupta and KJ Friston 2015; P Sterling and S Laughlin 2015). The contribution of this review is simply to show that these relationships can be united into a common theoretical framework.

## Neurobiology

> *“… classical thermodynamics… is the only physical theory of universal content which I am convinced, that within the framework of applicability of its basic concepts, will never be overthrown.”*
>
> — – (A Einstein 1949, from JD Bekenstein 2001)

This section reviews thermodynamic relationships in systems neuroscience with a focus on information and energy. Beginning with neurons, moving to neural networks, and concluding at the level of the brain as a whole, I discuss the energetics of processes such as learning and memory, excitation and inhibition, and the production of noise in neurobiological systems.

The central role of energy in determining the activity of neurons exposes the close connection between information and thermodynamics at the level of the cell. For instance, the process of depolarization, which occurs as a transition to *E*_*max*_ from a resting state *E*_*min*_, clearly shows that cellular information content is correlated with energy levels. In this respect, the resemblance between ion concentration gradients in neurons and temperature gradients in thermodynamic demons (i.e., agents that use information from their surroundings to decrease their thermodynamic entropy) is not a coincidence – in order to acquire information, neurons must expend energy to establish proper membrane potentials. Recall that Landauer’s principle (JMR Parrondo et al. 2015; MB Plenio and V Vitelli 2001) places a lower bound on the quantity of energy released into the surroundings with the removal of information from a system. Thus, reestablishing membrane potentials after depolarization – the neuronal equivalent of resetting a demon’s memory – dissipates energy. Because Landauer’s principle applies to all levels of structure, and cells process large quantities of information, neurons use energy efficiently despite operating at several orders of magnitude above the nominal limit. Parameters including membrane area, spiking frequency, and axon length have all been optimized over the course of evolution to allow neurons to process information efficiently (P Sterling and S Laughlin 2015). Examining the energetics of information processing in neurons reinforces the notion that, while it is often convenient to imagine the neuron to be a simple binary element, these cells are intricate computational structures that process more than one bit of information.

Relationships between information and energy can also be seen at the level of neural networks. Attractor networks naturally stabilize by seeking energy minima, and the relative positions of basins of attraction define the geometry of an energy landscape (DJ Amit 1992). As a result, the transition into an active attractor state occurs as a transition into an information-energy maximum. These transitions correspond to the generation of informational entities such as memories, decisions, and perceptual events (ET Rolls 2012). In this way, the energy basins of attractor networks may be analogous to lower-level cellular and molecular energy gradients; a transition between any number of distinguishable energy levels follows the passage of a finite quantity of information. Since processing information requires the expenditure of energy, competitive network features also underscore the need to minimize unnecessary information processing. Lateral inhibition at this level may optimize thermodynamic efficiency by reducing metabolic expenses associated with networks responding less robustly to entering signals. Another interesting thermodynamic property of networks concerns macrostates: the functional states of large-scale neural networks rest emergently on the states of neuronal assemblies (R Yuste 2015). As a result, new computational properties may arise with the addition of new layers of network structure. Finally, the energetic cost of information has influenced network connectivity by imposing selective pressures to save energy by minimizing path length between network nodes (E Bullmore and O Sporns 2009).

Again, in accordance with Landauer’s principle, the displacement of information from any system releases energy into the surroundings (MB Plenio and V Vitelli 2001; TL Duncan and JS Semura 2004). This principle can be understood by imagining an idealized memory device, such as the brain of a thermodynamic demon. Since information is conserved (L Susskind and G Hrabovsky 2014), and clearing a memory erases information, the thermodynamic entropy of the surroundings must increase when a demon refreshes its memory to update information. This fundamental connection between information, entropy, and energy appears in many areas of the neurobiology of learning. For example, adjusting a firing threshold in order to change the probability that a system will respond to a conditioned stimulus (Y Choe 2015; T Takeuchi et al. 2014) optimizes engram fitness by minimizing the quantity of energy needed for its activation (S Still et al. 2012). Recurrent collateral connections further increase engram efficiency by enabling a minimal nodal stimulus to elicit its full energetic activation (ET Rolls 2012). Experimental evidence also shows that restricting synaptic energy supply impairs the formation of stable engrams (JJ Harris et al. 2012). Because the formation and disassembly of engrams during learning and forgetting optimizes the growth and pruning of networks in response to external conditions, the process of learning is itself a mechanism for minimizing entropy in the brain (KJ Friston 2003).

As another example of a multiscale process integrated across many levels by thermodynamics, consider the active balance between excitation and inhibition in neurobiological systems. Maintaining proper membrane potentials and adequate concentrations of signaling molecules requires the expenditure of energy, so it is advantageous for systems of the brain to minimize the processing of unnecessary information – to “send only what is needed” (P Sterling and S Laughlin 2015). Balancing excitation and inhibition is therefore a crucial mechanism for saving energy. Theoretical evidence that this balancing maximizes the thermodynamic efficiency of processing Shannon information (B Sengupta et al. 2013) is consistent with experimental findings in several areas of research on inhibition. For instance, constant inhibitory modulation is needed to stabilize internal states, and hyperexcitation (e.g., in epilepsy, intoxication syndromes, or trauma) can decrease relative information by reducing levels of consciousness (B Haider et al. 2006; K Lehmann et al. 2012). Likewise, selective attention is mediated by the activation of inhibitory interneurons (G Houghton and SP Tipper 1996), and sensory inhibition appears to sharpen internal perceptual states (JS Isaacson and M Scanziani 2011). The need to balance excitation and inhibition at all levels of structure highlights the energetic cost of information.

A final example worth discussing is the relationship between thermodynamics and the production of noise in neurobiological systems. Noise is present in every system of the brain, and influences all aspects of the organ’s function (AA Faisal et al. 2008; ET Rolls and G Deco 2010; A Destexhe and M Rudolph-Lilith 2012). Even in the absence of any potential forms of classical stochastic resonance, the noise-driven exploration of different states may optimize thermodynamic efficiency by allowing a system to randomly sample different accessible configurations. Theoretical arguments suggest indeed that noise enables neural networks to respond more quickly to detected signals (ET Rolls 2012), and empirical evidence implicates noise as a beneficial means of optimizing the performance of diverse neurobiological processes (MD McDonnell and LM Ward 2011). For example, noise in the form of neuronal DNA breaking (JU Guo et al. 2011; K Herrup et al. 2013; P Tognini et al. 2015) could enhance plasticity, since any stochastically optimized configuration would be more likely to survive over time as, in this case, a strengthened connection in a modifiable network. Because noise is a form of relative entropy, optimizing the signal-to-noise ratio in any neurobiological system promotes the efficient use of energy.

At the level of the brain as a whole, the connection between information and thermodynamics is readily apparent in the organ’s functional reliance on energy (PJ Magistretti and I Allaman 2015), its seemingly disproportionate consumption of oxygen and energy substrates (e.g., ATP, glucose, ketones, etc.) (S Herculano-Houzel 2011; ME Raichle and DA Gusnard 2002), its vulnerability to hypoxic-ischemic damage (JP Dreier et al. 2013; PL Lutz et al. 2003) and in the reduction of consciousness often conferred by the onset of energy restrictions (RG Shulman et al. 2009; J Stender et al. 2016). All fMRI, PET, and EEG interpretation rests on the foundational assumption that changes in the information content of neurobiological systems can be inferred by observing energy changes (D Attwell and C Iadecola 2002; G Collell and J Fauquet 2015), and it is well known that the information processing capacities of neurobiological systems are limited by energy supply (C Howarth et al. 2012). Overall, these relationships are consistent with the form of information-energy equivalence predicted by Landauer’s principle and information-entropy bounds. The living brain appears to maintain a state of thermodynamic optimization.

## Consciousness and free will

> *“… science appears completely to lose from sight the large and general questions; but all the more splendid is the success when, groping in the thicket of special questions, we suddenly find a small opening that allows a hitherto undreamt of outlook on the whole.”*
>
> — – (L Boltzmann 1886, from HC Von Baeyer 1999)

Although neuroscience has yet to explain consciousness or free will at any satisfactory level of detail, relationships between information and energy seem to be recognizable even at this level of analysis. This section reviews attempts to conceptualize major properties of consciousness (unity, continuity, complexity, and self-awareness) as features of information processing in the brain, and concludes with a discussion of free will.

At any given moment, awareness is experienced as a unified whole. Physical information is the substrate of consciousness (A Annila 2016), and the law of conservation of information requires any minimal unit of information to be transferred into a thermodynamic system as a temporally unitary quantity. As a result, it is possible that the passage of perceptual time itself occurs secondarily to the transfer of information, and that the information present in any integrated system of the brain at any observed time is necessarily cohesive and temporally unified. In this framework, the passage of time would vary in proportion to a system’s rate of energy dissipation. Although it is possible that physical systems in general exchange information in temporally unitary quantities, it is likely that many of the familiar features of the perceptual unity of consciousness require the structure and activity of neural networks in the brain. The biological basis of this unity may be the active temporal consolidation of observed events by integrated higher-order networks (S Dehaene and JP Changeux 2011; SA Greenfield and TFT Collins 2005; A Revonsuo 1999; F Varela et al. 2001). An informational structure generated by the claustrum has been speculated to contribute to this experiential unity (FC Crick and C Koch 2005, MZ Koubeissi et al. 2014), but it has also been reported that complete unilateral resection of the system performed in patients with neoplastic lesions of the region produces no externally observable changes in subjective awareness (H Duffau et al. 2007). Overall, it appears unlikely that the presence of information in any isolated or compartmentalized network of the brain is responsible for generating the unified nature of conscious experience.

While perceptual time is likely the product of a collection of related informational processes rather than a single, globalized function mediated by any one specific system of the brain, some of the perceptual continuity of consciousness may result from the effectively continuous flow of thermodynamic information into and out of integrated systems of the brain. In this framework, the quantum (M Prokopenko et al. 2014) of perceptual time would be the minimal acquisition of information, and the entrance of information into neurobiological systems would occur alongside the entrance of energy. This relationship is implicit in the simple observation that the transition of a large-scale attractor network is progressively less discrete and smoother in time than the activation of a small-scale engram, the propagation of a cellular potential, the docking of a vesicle, the release of an ion, and so forth. Likewise, electroencephalography shows that the summation of a large number of discrete cellular potentials can accumulate into an effectively continuous wave as a network field potential (PL Nunez and R Srinivasan 2006), disruptions of which are often correlated with decreases in levels of consciousness (H Blumenfeld and J Taylor 2003). It is also well known that higher frequency network oscillations tend to indicate states of wakefulness and active awareness, while lower frequency oscillations tend to be associated with internal states of lesser passage of perceptual time, such as dreamless sleep or unconsciousness. The possibility that the experiential arrow of time and the thermodynamic arrow of time share a common origin in the flow of information is supported both by general models of time in neuroscience and the physical interpretation of time as an entropy gradient (L Mlodinow and TA Brun 2014; OC Stoica 2008).

The subjective complexity of consciousness may show that extensive network integration is needed for maximizing the mutual thermodynamic information and internal energy content of systems of the brain (JS Torday and WB Miller Jr 2016). An exemplary structure enabling such experience, likely one of many that together account for the subjective complexity of consciousness, is the thalamocortical complex (Y Hannawi et al. 2015; RS Calabrò et al. 2015). The functional architecture of such a network may show that, at any given moment in the internal model of a living brain, a wide range of integrated systems are sharing mutual sources of thermodynamic information. This pattern of structure may reveal that the perceptual depth and complexity of conscious experience is a direct product of recognizable features of the physical brain. However, it also seems that extensive local cortical processing of information is necessary for producing a refined and coherent sensorium within a system, and that both the thalamocortical complex and the brain stem are involved in generating the subjective complexity of consciousness (GM Edelman et al. 2011; LM Ward 2011). The dynamics of attractor networks at higher levels of network structure may show that quantities of complex internal information can be observed as changes in cortical energy landscapes (ET Rolls 2012), with a transition between attractor states following the transfer of information. The degree of subjective complexity of information enclosed by such a transition would be proportional to the degree of structural integration of underlying networks.

Self-awareness likely arose as a survival necessity rather than as an accident of evolution (F Fabbro et al. 2015), and rudimentary forms of self-awareness likely began to appear early in the course of brain evolution as various forms of perceptual self-environment separation. As a simple example, consider the tickle response (DJ Linden 2007), which requires the ability to differentiate self-produced tactile sensations from those produced by external systems. The early need to distinguish between self-produced tactile states and those produced by more threatening non-self sources may be reflected by the observation that this recognition process is mediated to a great extent by the cerebellum (SJ Blakemore et al. 2000). While it is possible that other similar developments began occurring very early on, the evolutionary acquisition of the refined syntactical and conceptual self present in the modern brain likely required the merging of pre-existing self networks with higher-level cortical systems. The eventual integration of language and self-awareness would have been advantageous for coordinating social groups (MS Graziano 2013), since experiencing self-referential thought as inner speech facilitates verbal communication. Likewise, the coupling of self-awareness to internal sensory, cognitive, and motor states (T Metzinger 2004; G Northoff et al. 2006) may be advantageous for maximizing information between systems within an individual brain. Neuropsychological conditions involving different forms of agnosia, neglect, and self-awareness deficits do show that a reduced awareness of self-ownership of motor skills, body parts, or perceptual states can result in significant disability (F Fabbro et al. 2015; S Chokron et al. 2016; MD Orfei et al. 2007; M Overgaard 2011; A Parton et al. 2004; M Tsakiris 2010; A Morin 2006; GP Prigatano 2009). Since experiencing self-awareness optimizes levels of mutual information between the external world and the brain’s internal model (MA Apps and M Tsakiris 2014), and this activity decreases thermodynamic entropy (JS Torday and WB Miller Jr 2016), self-awareness may be a mechanism for optimizing the brain’s consumption of energy.

Thermodynamic information is also interesting to consider in the context of free will. The brain is predictable within reason, and the performance of an action can be predicted before a decision is reported to have been made (P Haggard 2008). Entities such as ideas, feelings, and beliefs seem to exist as effectively deterministic evaluations of information processed in the brain. Whether or not the flow of information is subject to the brain’s volitional alteration, neuroscience also shows that information can be internally real to a system of the brain, even if this information is inconsistent with an external reality. That the brain can generate an externally inconsistent internal reality is demonstrated by phenomena such as confabulation, agnosia, blindsight, neglect, commissurotomy and hemispherectomy effects, placebo and nocebo effects, reality monitoring deficits, hallucinations, prediction errors, the suspension of disbelief during dreaming, the function of communication in minimizing divergence between internal realities, the quality of many kinds of realistic drug-induced experiences, and the effects of many neuropsychological conditions. The apparent fact that subjective reality is an active construction of the physical brain has even led to the proposal of model-dependent realism (SW Hawking and L Mlodinow 2011) as a philosophical paradigm in the search for a unified theory of physics. In any case, it is likely that beliefs, including those in free will, exist as information, and that their internal reality is a restatement of its frequently observer-dependent nature.

## Empirical outlook

Before concluding, it is worth reviewing a few notable experiments in greater detail. While considerable advances have been made in discovering how neurobiological systems operate according to principles of thermodynamic efficiency (S Laughlin and P Sterling 2015), relationships between information and energy in the brain are only beginning to be understood. The following studies are examples of elegant and insightful experiments that should inspire future research.

Several recent brain imaging studies support the proposal (A Annila 2016) that thermodynamics is able to explain a number of mysteries involving consciousness. For example, J Stender et al. 2016 used PET to measure global resting state energy consumption in 131 brain injury patients with impairments of consciousness as defined by the revised Coma Recovery Scale (CRS-R). The preservation of consciousness was found to require a minimal global metabolic rate of ≈ 40% of the average rate of controls; global energy consumption above this level was reported to predict the presence or recovery of consciousness with over 90% sensitivity. These results must be replicated and studied in closer detail before their specific theoretical implications are clear, but it is now established that levels of consciousness are correlated with energetic metrics of brain activity. To what extent there exists a well-defined “minimal energetic requirement for the presence of conscious awareness” (J Stender et al. 2016) remains an open question. However, the empirical confirmation of a connection between consciousness and thermodynamics introduces the possibility of developing new experimental methods in consciousness research.

Neurobiological systems, and biological systems in general (HC Von Baeyer 1999; ED Schneider and D Sagan 2005), can be considered thermodynamic demons in the sense that they are agents using information to decrease their thermodynamic entropy. Landauer’s principle requires that, in order not to violate any known laws of thermodynamics, such agents dissipate heat when erasing information from their memory storage devices. In an experimental test of this principle, reviewed along with similar experiments in JMR Parrondo et al. 2015, A Bérut et al. 2012 studied heat dissipation in a simple memory device created by placing a glass bead in an optical double-well potential. Intuitively, this memory stored a bit of information by retaining the bead on one side of the potential rather than on the alternative. By manipulating the height of the optical barrier between wells, researchers moved the bead to one side of the memory without determining its previous location in the potential. This process was therefore logically irreversible, requiring the erasure of prior information from the memory device. Landauer’s principle predicts that, since information is conserved, the entropy of the memory’s surroundings must increase when this occurs. A Bérut et al. 2012 have verified that energy is emitted when a memory is cleared. As noted by the authors, “this limit is independent of the actual device, circuit or material used to implement the irreversible operation.” It would be interesting to study the erasure principle in the context of neuroscience.

Experimental applications of information theory in cell biology have already led to the discovery of general principles of brain organization related to thermodynamics (S Laughlin and P Sterling 2015). In one particularly interesting study, JE Niven et al. 2007 measured the energetic efficiency of information coding in retinal neurons. Intracellular recordings of membrane potential and input resistance were used to calculate rates of ATP consumption in response to different background light intensities. These rates of energy consumption were then compared with rates of Shannon information transmission in order to determine metabolic performance. It was found that metabolic demands increase nonlinearly with respect to increases in information processing rate: thermodynamics appears to impose a “law of diminishing returns” on systems of the brain. The authors interpret these results as evidence that nature has selected for neurons that minimize unnecessary information processing. Studying how thermodynamics has influenced cellular parameters over the course of evolution is likely to raise many new empirically addressable questions.

## Conclusion

This article has reviewed information-energy relationships in the hope that they may eventually provide a general framework for uniting theory and experiment in neuroscience. The physical nature of information and its status as a finite, measurable resource are emphasized to connect neurobiology and thermodynamics. As a scientific paradigm, the information movement currently underway in physics promises profound advances in our understanding of the relationship between energy, information, and the physical brain.

## Acknowledgements

I am grateful to Baroness Susan Greenfield, Dr. Francesco Fermani, Dr. Karl Friston, Dr. Biswa Sengupta, Dr. Roy Frieden, Dr. Bernard Baars, Dr. Brett Clementz, Dr. Cristi Stoica, Dr. Satoru Suzuki, Dr. Paul King, Guillem Collell Talleda, Dr. Jordi Fauquet, and others who have helped me improve these ideas. I am also grateful to Dr. Shanta Dhar and her team for introducing me to academic research, to Jim Reid for introducing me to biology, to Alex Tisch for introducing me to physics, and to those affiliated with the Department of Neurosurgery at the University of Virginia Medical Center for introducing me to neuroscience.

